# Performance evaluation of biosurfactant stabilized microbubbles in enhanced oil recovery

**DOI:** 10.1101/504431

**Authors:** Gunaseelan Dhanarajan, Shaheen Perveen, Anirban Roy, Sirshendu De, Ramkrishna Sen

## Abstract

Microbubble technology is increasingly finding applications in biomedical engineering, soil remediation and wastewater treatment. Recently, the use of surfactant microbubbles has been studied as an alternate to aqueous solution in pollutant remediation since they have the advantage of improving the contact with the contaminant due to their surface properties. In this research endeavor, the application potential of microbubble suspension generated using a lipopeptide biosurfactant produced by a marine strain of Bacillus megaterium in tertiary oil recovery was investigated. The microbubbles were generated using a high speed homogenizer and their properties such as stability and size distribution were studied. The microbubble suspension was used as flooding agent to recover gear oil from an artificially saturated sand packed column. The performance of microbubbles in tertiary oil recovery was compared with that of aqueous biosurfactant solution. It was found that microbubble suspension generated using biosurfactant had higher oil recovery efficiency (46%) than aqueous solution (36%).

Moreover, the pressure buildup across the sand packed column was fairly low while using microbubble suspension. The increased oil recovery using microbubbles can be attributed to their effective permeation through the pores of sand packed column and closer contact between biosurfactant molecules and oil. Thus, the results obtained in this study convincingly indicate that biosurfactant stabilized microbubble suspension, due to its higher performance and lower injection pressure requirement, can serve as a potentially efficient flooding agent for tertiary oil recovery.

## 1. INTRODUCTION

Present oil recovery operations mainly rely on natural reservoir pressure and gas/water flooding into the reservoir to drive the oil to producing wells. These technologies can result in a maximum recovery of 40–45% of the original oil in place and thus, leaving two-thirds of the oil in the reservoir (Sen, 2008). However, the impending petroleum energy crisis worldwide has necessitated the development of enhanced oil recovery (EOR) techniques as they provide an additional chance to extract about 20–25% more oil from the mature oil wells (Sen, 2008). A recent survey by Grand View Research revealed that worldwide EOR market volume in 2013 was 2.68 billion barrels, dominated by North America which accounts for 36.4% of the total market. The EOR market volume is expected to grow at a CAGR of 29.9% to reach 16 billion barrels by 2020. Correspondingly, the worldwide EOR market is expected to reach $283 billion by 2020 (https://www.grandviewresearch.com/press-release/global-enhanced-oil-recovery-eor-market; accessed on 30 September 2016). The EOR process mainly involves the application of surfactants to mobilize the oil left in the mature oil fields as they reduce the interfacial tension between the brine and oil and increase the sweep efficiency. However, the commonly-used synthetic surfactants are often toxic to the environment, especially when present with oil (Patel et al., 2015). This calls for the use of microbially derived products such as biosurfactants and biopolymers in place of chemically synthesized counterparts. The features that make them promising alternatives to synthetic surfactants for EOR include lower toxicity, higher biodegradability, superior surface active and emulsification properties, and stability at extremes of temperature, pH and salinity (Dhanarajan et al., 2015; Mukherjee et al., 2006; Rangarajan et al., 2012). A state-of-the-art technology that may be more efficient in EOR is the application of biosurfactants in the form of microbubble suspension. The microbubbles are encapsulated with a thin film of multilayer biosurfactant molecules and are comprised of about 60–70% gas by volume and therefore, form a low density fluid. Thus, the microbubbles can pass through the pores of reservoir effectively and improve oil recovery efficiency. The use of microbubble suspension provides additional advantages such as lower pressure build-up and lesser requirement of biosurfactant (Hashim et al., 2012). Furthermore, microbubbles are reported to be stable at elevated pressure and temperature (Shivhare and Kuru, 2014). Application of microbubble suspension generated using synthetic surfactants was found to be more efficient in flushing different hydrophobic organic contaminants and heavy metals from soil in comparison to aqueous surfactant solutions on the basis of pollutant removed per mole of surfactant used (Boonamnuayvitaya et al., 2009; Roy et al., 1995). These applications exploit the properties of microbubbles such as large interfacial area and enhanced mass transfer ability. However, studies on EOR using surfactant microbubble suspension are scarce. Thus, the present study was aimed at employing microbubble suspension synthesized using an ecofriendly lipopeptide biosurfactant for EOR in a sand packed column.

## 2. MATERIALS AND METHODS

### 2.1. Production and isolation of biosurfactant

Biosurfactant was produced using a marine Bacillus megaterium (isolated from Andaman and Nicobar Islands, India) in noodle processed water supplemented with chemical fertilizers as described previously (Dhanarajan et al., 2014). After 48 h of fermentation, the culture broth was centrifuged at 10000 rpm for 10 min to separate the cells. The cell free supernatant was acidified to pH 2.0 using 6N HCl to precipitate the biosurfactant. The precipitate was centrifuged and the pellet was lyophilized to get the biosurfactant in crude form. The concentration of crude biosurfactant produced by the marine bacterium was measured to be 6.5 g L^-1^. The biosurfactant product was found to contain a mixture of iturin (23%), fengycin (44%) and surfactin (33%) type of lipopeptides (Dhanarajan et al., 2016). Although, surfactin is reported to be the most potent surface active agent among the lipopeptide biosurfactants, due to the cost-intensive downstream processing procedures to separate surfactin from lipopeptide mixture, the crude mixture was used as such in this study (Mukherjee et al., 2006)

### 2.2. Determination of emulsification activity

Emulsification activity of the lipopeptide biosurfactant was tested with gear oil, gasoline, diesel and hexadecane. Equal volume of hydrocarbon and 500 mg L^-1^ biosurfactant solution were added in a centrifuge tube and mixed vigorously using a vortex mixer for 5 min and incubated for 24 h without any disturbance. The emulsification index (E24) was determined as the percentage height of emulsified layer upon the total height of the mixture.

### 2.3. Influence of lipopeptide concentration on surface tension and interfacial tension between oil and water

Biosurfactant solutions of varying lipopeptide concentrations (25 – 1000 mg L^-1^) were prepared. The pH of all samples was adjusted to 8 before measurement. The surface and interfacial tension of the solutions was determined using tensiometer by Du Noüy ring method using a tensiometer (Lauda TD 1C, Germany).

### 2.4. Generation and characterization of microbubble suspension

Microbubbles were synthesized by two methods namely, high speed homogenization and membrane emulsification. Firstly, a high speed homogenizer (Model: D500; Make: Scilogex, USA) was used (Fig. 1a). The homogenizer shaft was partially immersed into biosurfactant solution (250 and 500 mg L^-1^) and the sample was homogenized at 10,000 rpm for 3 minutes. Then, the suspension was left standing for a few seconds until the microbubbles moved to top phase. The milky white microbubble suspension was separated to analyze for size distribution and stability. About five microliters of microbubble sample was pipetted out and placed on a concavity slide (Himedia, India) and observed under Nikon Eclipse 50i microscope (Nikon Corporation, Japan) with a high resolution camera (Nikon Digital Sight DS-Fi1) attached to it. Nikon Elements Software-D was used for capturing all microbubble images. The images were examined using ImageJ software (National Institutes of Health, Bethesda, Maryland, USA; version 1.46r) and statistically analyzed using Origin Pro software (Version 8.5). Size distribution was determined by measuring the sizes of at least 100 microbubbles from recorded images. To measure the stability of microbubbles, about 5 mL of microbubble suspension was taken in a separate vial, stored at 28 °C and the time span for which microbubble stay before collapsing into liquid was recorded. Gas hold-up, the volumetric fraction of gas present in the suspension, was estimated using the following equation.

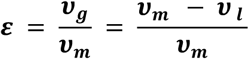

where, ***ε*** is gas hold-up, ***ν***_***g***_ is volume of gas, ***ν***_***m***_ is volume of microbubble suspension, ***v***_***l***_ is volume of liquid drained from microbubble suspension after collapse. Alternatively, synthesis of near mono-dispersed microbubbles was attempted by membrane emulsification method using a hollow fiber membrane of pore size approximately 44 kDa as reported elsewhere (Kukizaki, 2009). The hollow fiber membrane was synthesized in our laboratory using polysulfone and polyethylene glycol (PEG) (molecular weight 200). A polymer blend solution was prepared (PSf:PEG) by dissolving them in ratio (wt%) of 18:3 in di-methyl formamide DMF with slow stirring at 60 °C (Roy et al., 2016a). The polymer solution was then used to prepare hollow fiber membrane with a pore size of approximately 44 kDa (Roy et al., 2016b). The solution was then used to spin hollow fiber membranes. The authors have resorted to an in house technology as reported by Thakur and De using syringe-in-syringe assembly (Thakur and De, 2012). Two co-axial syringes have been used to extrude membranes which were further characterized for its pore size. Once spun, the membrane was kept in distilled water overnight to complete phase inversion and minimize presence of solvent in the membrane matrix. Then, 50 fibers were cut into 8 inches long pieces and potted into ½ inch PVC pipe (1 inch length) in a u-shape so that the membranes protrude from it and are exposed to water (Figure S1). Compressed air was passed through the membrane pores into the lipopeptide solution to generate microbubbles. However, the process of microbubble generation was found to be very slow in comparison to homogenization method. Hence, oil recovery experiments were carried out using microbubble suspension generated by high speed homogenization

**Fig. 1.**
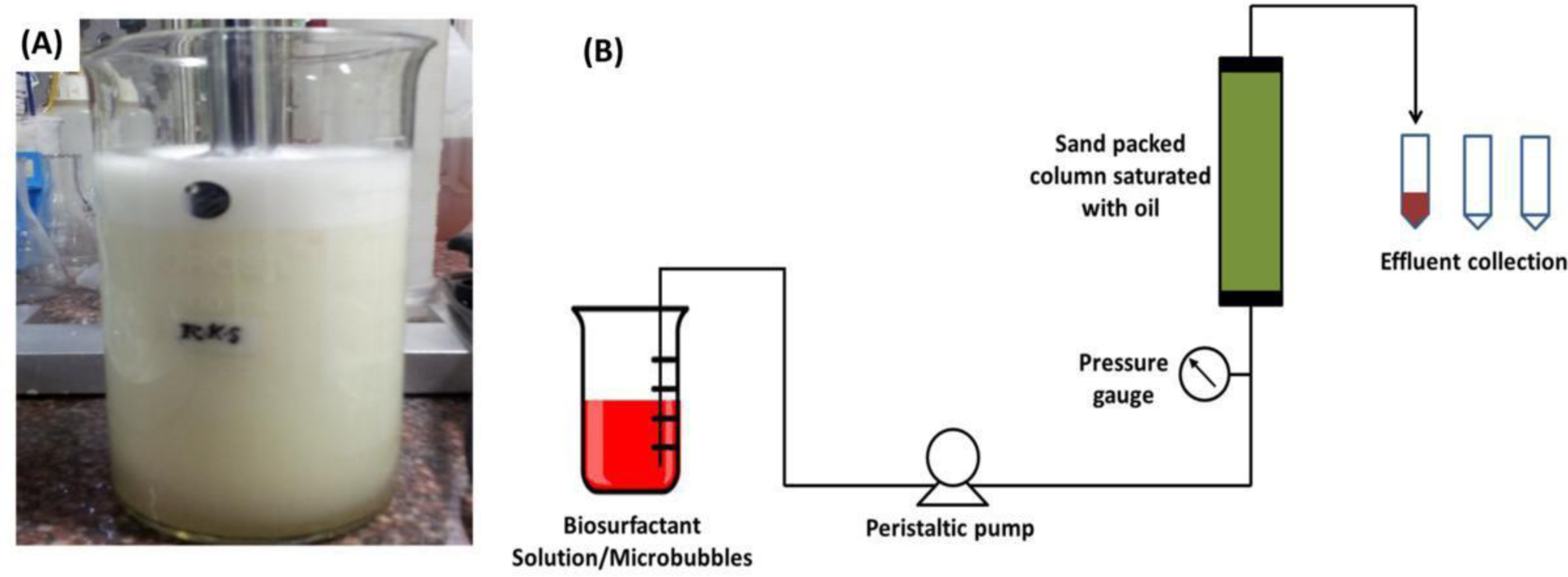
(a) Microbubble suspension generated using biosurfactant solution. (b) Scheme of the experimental set-up used for oil recovery studies.

### 2.5. Oil recovery studies in sand packed column

To perform the tertiary oil recovery studies, a 4 × 10 cm column was packed with approximately 290 g of sand (20–30 mesh). The column was initially flooded with deionized water at a pressure gradient close to zero to determine the pore volume as ∼ 62 mL. The column was saturated with 80W gear oil and then brine solution (5% NaCl) was passed until no more oil discharged from the column, which brought the column to a condition of residual oil saturation. The volume of oil saturated (oil-in-place) in the sand packed column was estimated by measuring the difference between the initial volume injected into the column and the volume recovered with brine flooding. After flushing with brine, six pore volumes of biosurfactant solution at different concentrations was injected into the column at a rate of 5 mL min^-1^, followed by two pore volumes of water. The liquid discharging from the column was collected and the volume of oil recovered was recorded. For microbubble flooding, microbubble suspension generated using the same volume of biosurfactant solution was pumped at the aforementioned flow rate. Scheme of the experimental set-up used in this study is given in Fig.1b.

## 3. RESULTS AND DISCUSSION

### 3.1. Emulsification activity of the biosurfactant

The lipopeptide biosurfactant produced by the marine B. megaterium showed stable emulsification activity against various hydrocarbons as shown in Fig. 2. Among the tested hydrocarbons, maximum E24 was exhibited against diesel (61%), while minimum was obtained with engine oil (44%). The E24 values obtained in this study were found to be comparable with that in previous literature reports. A lipopeptide biosurfactant synthesized by Pseudomonas nitroreducens showed emulsification activity of 40–60% for various hydrocarbons and oils (de Sousa and Bhosle, 2012), whereas that from Bacillus subtilis K1 exhibited in the range of 30–55% (Pathak and Keharia, 2014). Thus, the results indicated that the biosurfactant could be a potential agent in enhanced oil recovery for it showed stable emulsification activity with both aliphatic and aromatic hydrocarbons.

**Fig. 2.**
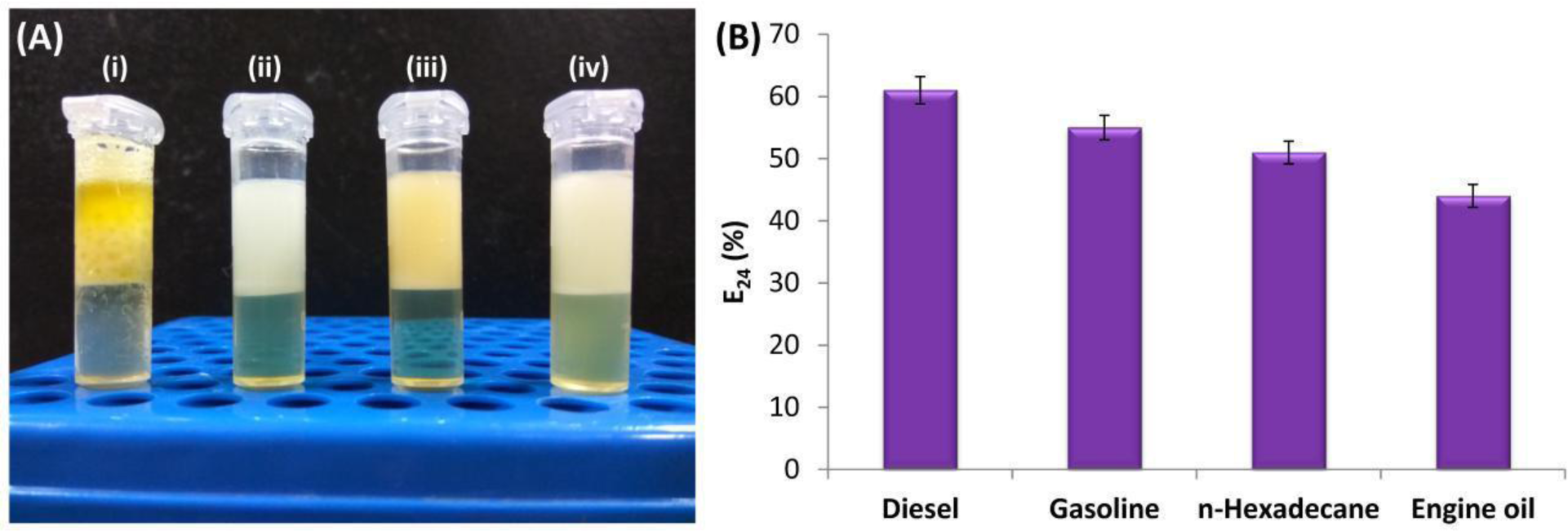
(a) Photographs of emulsions formed with biosurfactant and (i) gear oil, (ii) n-hexadecane, (iii) gasoline and (iv) diesel. (b) Emulsification index (E24%) of biosurfactant against various hydrocarbons

### 3.2. Surface and interfacial tension measurements

The CMC of crude lipopeptide biosurfactant in water at pH 8 was found to be 250 mg L^-1^ (Fig 3a), giving a minimum surface tension of 28.25 mN m^-1^. The biosurfactant reduced the interfacial tension between water to oil from 18.6 to 1.5 mN m^-1^ (Fig. 3b). The CMC of the crude biosurfactant mixture produced by the marine strain was similar to the reported values elsewhere. Al-Wahaibi et al. observed that surfactin lipopeptide decreased the interfacial tension between water and Arabian crude oil from 22.84 to 3.79 mN m^-1^ at a CMC of 300 mg L^-1^ (Al-Wahaibi et al., 2014). Similarly, lipopeptide biosurfactant produced by a B. subtilis strain isolated from Brazilian crude oils reduced the IFT between water and oil from 22 to 1 mN m^-1^ at a concentration of 200 mg L^-1^ (Pereira et al, 2013).

**Fig. 3.**
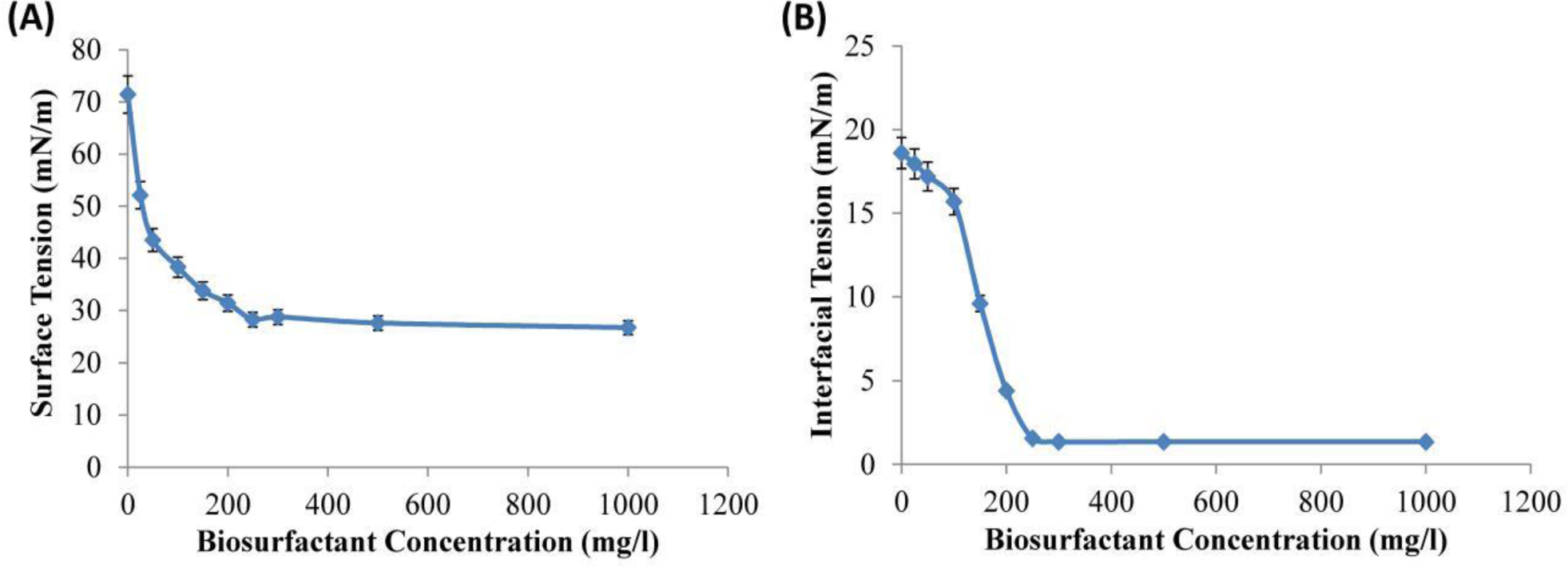
(a) Surface tension values of deionized water against various concentrations of biosurfactant. (b) Interfacial tension values between water and oil at different concentrations of biosurfactant.

### 3.3. Characterization of biosurfactant stabilized microbubbles

Between the two tested methods, high speed homogenization resulted in formation of larger volume of microbubbles at a faster rate as opposed to membrane emulsification. Hence, further studies were carried out using microbubbles synthesized by homogenizer. The change in concentration of biosurfactant was found to affect the size distribution and stability of microbubbles. Figure 4 shows the optical micrographs of microbubbles synthesized by homogenizer using various concentrations of biosurfactant solution. A rightward shift in the size distribution of microbubbles was observed as the concentration of biosurfactant solution was increased, indicating that larger bubbles were formed at higher concentration of biosurfactant molecules. The mean diameter of microbubbles synthesized using 250 mg L^-1^ biosurfactant solution was measured to be 61μm. Whereas, increase in concentration to 500 mg L^-1^ resulted in formation of larger microbubbles with a mean diameter of 72μm. Similarly, the stability of microbubble suspension improved as the concentration of biosurfactant solution was increased. The lifetime of the suspensions synthesized using 250 and 500 mg L^-1^ biosurfactant concentrations was 16 h and 18.5 h, respectively. The longer lifetime of microbubbles might be due to increased number of lipopeptide molecules on an individual microbubble at higher biosurfactant concentration. This can be validated by taking the gas hold-up into account. The gas hold-up values of microbubble suspensions synthesized using 250 and 500 mg L^-1^ biosurfactant solutions were 0.68 and 0.6, respectively. The lower gas hold-up value at higher biosurfactant concentration indicates that the fraction of liquid around the microbubble was more, which in turn increases the number of lipopeptide molecules on each microbubble. Thus, the stability of the suspension was higher in case of microbubbles generated using highly concentrated biosurfactant solution.

**Fig. 4.**
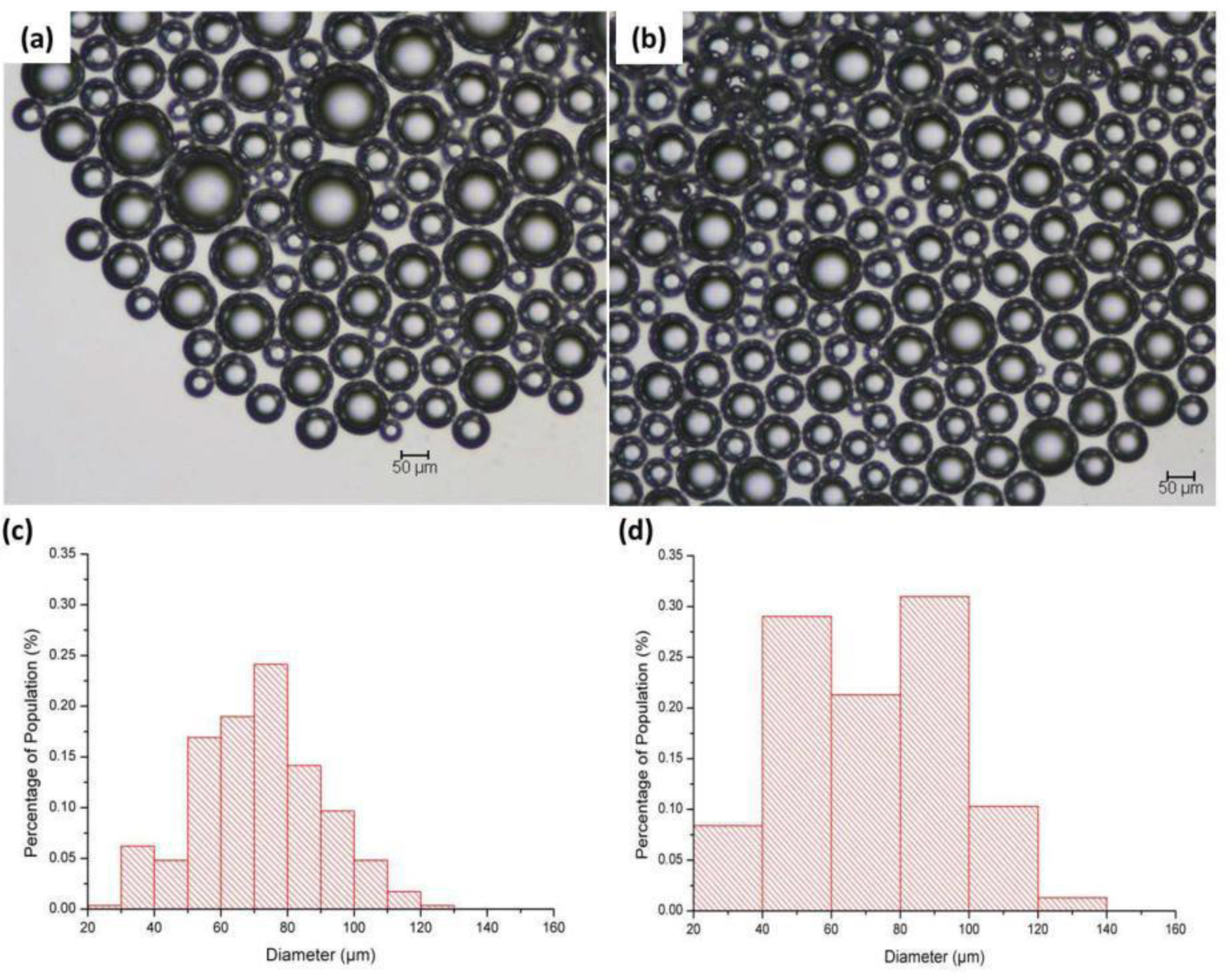
Optical micrographs and size distribution of microbubbles generated using 250 mg/L (a & c) and 500 mg/L (b & d) biosurfactant solutions, respectively.

### 3.4. Enhanced oil recovery using biosurfactant solution and microbubbles

The effect of biosurfactant flooding in the form of aqueous solution and microbubble suspension on oil recovery from a sand packed column is given in Fig. 5. As shown in the figure, oil recovery efficiency was 31 ± 1.5% and 36 ± 1.9% when flooding with six pore volumes of 250 and 500 mg L^-1^ biosurfactant solutions, respectively. However, the volumes of microbubble suspensions generated using the same biosurfactant solutions were 240 mL and 295 mL, which correspond to approximately four and five pore volumes of the column, respectively. Flooding with such microbubble suspensions resulted in oil recovery of 29 ± 1.4% and 46 ± 2.2%, respectively. The recovery of oil using biosurfactant molecules might be attributed to mechanisms such as oil displacement from pores, solubilization in lipopeptide micelles, oil dispersion due to electric repulsion between oil and soil particles (Tsakiroglou et al., 2013). Simultaneous flooding of biosurfactant and gas as microbubble suspension reduces the fluid mobility, which enhances fluid residence time and the contact efficiency between the lipopeptide and oil. This might have improved the recovery efficiency of the biosurfactant in microbubble form as compared to aqueous solution. The surface area of the microbubbles might also have played a significant role in the enhanced recovery of oil from the sand packed column (Lim et al., 2015). Furthermore, due to the low density, the microbubbles can pass through the pores of sand packed column effectively and improve oil recovery. Thus, the recovery efficiency was more in case of microbubble flooding in comparison to aqueous solution on the basis of oil recovered per gram of biosurfactant used at 500 mg L^-1^. However, similar oil recovery efficiencies were observed using the microbubble suspension and aqueous solution at lower biosurfactant concentration. This behaviour could be attributed to the critical micelle concentration of the crude biosurfactant, which is 250 mg L^-1^. At CMC, the stability of microbubbles was lower and thus bubble dissolution was observed in the feed tank during flooding. Hence, lower oil recovery% was achieved for microbubble suspension at 250 mg L^-1^ biosurfactant. Moreover, it has been reported that surfactant concentrations higher than the CMC are required to achieve the maximum effect of the surfactant to stabilize microbubbles (Xu et al., 2009). On the other hand, above CMC, stable biosurfactant micelles might have enhanced microbubble stability and oil solubilization. This shows that the concentration of biosurfactant need be more than the CMC to generate microbubbles for EOR applications. Even though the microbubble suspension was stable for a period of about 16 h in ambient conditions for 250 mg L^-1^ biosurfactant, flooding into the sand packed column resulted in coalescence and collapse of a fraction of microbubbles as observed at the column exit. This might be due to various factors such as the pore structure, flooding rate and the type of medium. On the other hand, bubble coalescence was relatively less for microbubble suspension synthesized using 500 mg L^-1^ biosurfactant. However, the porosity and the permeability are expected to be much lesser in oil reservoirs, which might impede the flooding of microbubbles. Thus, it is important to study the feasibility of the microbubble

**Fig. 5.**
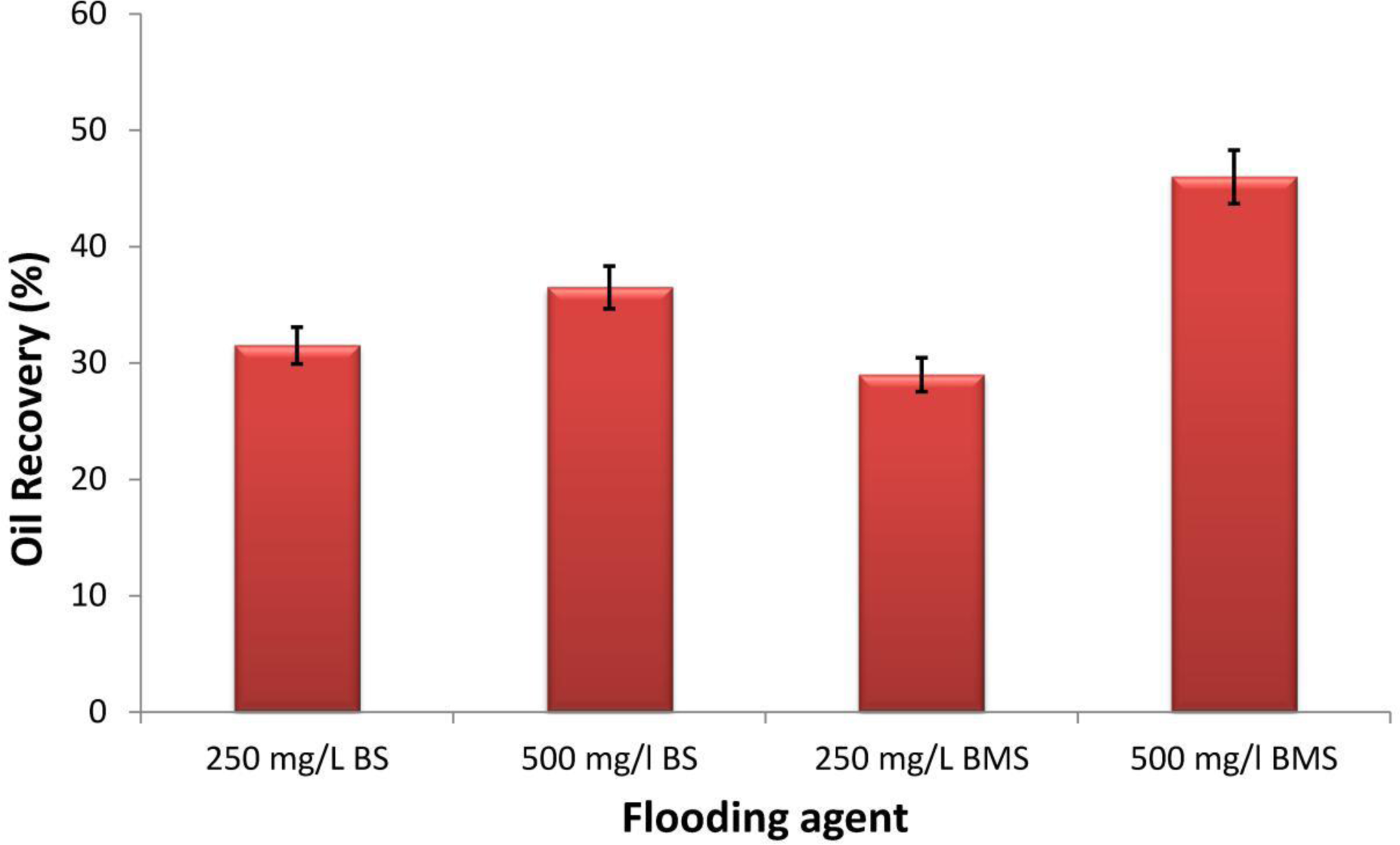
Tertiary oil recovery in a sand packed column using biosurfactant solution (BS) and microbubble suspension (BMS)

### 3.5. Pressure buildup in sand packed column

Application of surfactant or polymer solution generally results in pore plugging in the reservoirs, which might cause excessive pressure build**-**up. Hence, it is a critical factor that influences the use of surfactant solutions for EOR in situ. The pressure build-up in the sand packed column while flooding with biosurfactant solution and microbubble suspension is shown in Fig. 6. The trend in variation of pressure was found to be similar in all cases. As shown in the figure, the pressure build-up across the column increased sharply for the first two volumes and gradually thereafter. However, the pressure build-up in case of biosurfactant solution was observed to be higher as compared to microbubble suspension. Flushing with 500 mg L^-1^ biosurfactant solution increased the pressure up to 82 kPa, whereas microbubble flooding resulted in relatively low pressure build-up of 40 kPa. On the other hand, when the flushing solution was switched to water, the pressure decreased in case of biosurfactant solution and increased for microbubble flooding. As the biosurfactant flooding was performed in up-flow mode, microbubbles, owing to buoyancy and low density, tend to move up effectively through the sand packed column. Thus, the pressure build-up was found to be lower in case of microbubble flooding. The lower injection pressure requirement during microbubble flooding makes this process advantageous over aqueous solution for in situ EOR applications.

## 4. CONCLUSION

In the present study, microbubble suspension generated using a microbially derived lipopeptide biosurfactant was employed for enhanced oil recovery and the performance was compared with aqueous biosurfactant solution. The microbubble suspension was found to be stable for more than 16 h and the size of the bubbles was ranging from 20–120μm in diameter. Microbubble suspension generated from 500 mg L^-1^ biosurfactant solution recovered 46% of oil that remained in sand packed column as compared to 34% by aqueous solution at the same biosurfactant concentration. However, the injection pressure required for biosurfactant flooding was significantly lower in case of microbubble suspension in comparison to aqueous solution. Thus, this study indicates the potential of biosurfactant flooding in the form of microbubble suspension for in situ EOR applications due to its higher recovery efficiency and better injectivity.

## 5. ACKNOWLEDGEMENTS

The authors thankfully acknowledge the financial support received from the Department of Biotechnology, Government of India, for the project grant (No.: BT/PR6909/PBD/26/391/2013, 21/03/2014).

---

**Table 1.** Comparison of percentages of oil recovery using lipopeptide biosurfactants

| Microorganism | Biosurfactant<br>Concentration (g L<sup>-1</sup>) | Flooding solution | Oil<br>Recovery% | Reference |
| --- | --- | --- | --- | --- |
| *B. subtilis* | 0.1 | Aqueous biosurfactant<br>solution | 22 | Pereira et<br>al, 2013 |
| *B. subtilis* B30 | 0.5 | Aqueous biosurfactant<br>solution | 31 | Al-Wahaibi,<br>2014 |
| *Bacillus*<br>*subtilis* BS-37 | 0.3 | Aqueous biosurfactant<br>solution | 37 | Liu et al.,<br>2015 |
| *B. atrophaeus*5-2a | 0.77 | Aqueous biosurfactant<br>solution | 22 | Zhang et al.,<br>2016 |
| *Bacillus* Sp. | 0.5 | Aqueous biosurfactant<br>solution | 34 | Joshi and<br>Desai, 2013 |
| *B. megaterium* | 0.25 | Aqueous biosurfactant<br>solution | 31 | This study |
| *B. megaterium* | 0.5 | Aqueous biosurfactant<br>solution | 36 | This study |
| *B. megaterium* | 0.25 | Biosurfactant stabilized<br>microbubbles | 29 | This study |
| *B. megaterium* | 0.5 | Biosurfactant stabilized<br>microbubbles | 46 | This study |

## Figure Captions

**Fig. 6.** Pressure build-up across the sand packed column while flushing with biosurfactant solution (BS) and microbubble suspension (BMS). Arrow indicates change of flooding solution from BS to brine.

